# Anionic lipid catalyzes the generation of cytotoxic insulin oligomers

**DOI:** 10.1101/2025.01.14.633028

**Authors:** Jhinuk Saha, Audrey Wolszczak, Navneet Kaur, Malitha C. Dickwella Widanage, Samuel D McCalpin, Riqiang Fu, Jamel Ali, Ayyalusamy Ramamoorthy

## Abstract

Misfolding and aggregation of proteins into amyloidogenic assemblies are key features of several metabolic and neurodegenerative diseases. Human insulin has long been known to form amyloid fibrils under various conditions, which affects its bioavailability and function. Clinically, insulin aggregation at recurrent injection sites poses a challenge for diabetic patients who rely on insulin therapy. Furthermore, decreased responsiveness to insulin in type 2 diabetic (T2D) patients may lead to its overproduction and accumulation as aggregates. Earlier reports have reported that various factors such as pH, temperature, agitation, and the presence of lipids or other proteins influence insulin aggregation. Our present study aims to elucidate the effects of non-micellar anionic DMPG (1,2-dimyristoyl-sn-glycero-3-phosphoglycerol) lipids on insulin aggregation. Distinct pathways of insulin aggregation and intermediate formation were observed in the presence of DMPG using a ThT fluorescence assay. The formation of soluble intermediates, alongside large insulin fibrils, was observed in insulin incubated with DMPG via TEM, DLS and NMR, as opposed to insulin aggregates generated without lipids. ^13^C magic angle spinning solid-state NMR and FTIR experiments indicated that lipids do not alter the conformation of insulin fibrils but do alter the time scale of motion of aromatic and aliphatic sidechains. Furthermore, the soluble intermediates were found to be more cytotoxic as compared to fibrils generated with or without lipids. Overall, our study elucidates the importance of anionic lipids in dictating the pathways and intermediates associated with insulin aggregation. These findings could be useful in determining various approaches to avoid toxicity and enhance the effectiveness of insulin in therapeutic applications.

## Introduction

Amyloid aggregation has evolved as a key underlying mechanism of several human diseases. For instance, diseases like Type-2 diabetes (T2D), Parkinson’s disease (PD), Alzheimer’s disease (AD), Huntington’s Disease, and Amyotrophic lateral sclerosis (ALS) involve deposition of amyloid aggregates of one or more proteins within their pathophysiology^1–6^. Amyloid aggregation and insulin ball formation under the skin has also been observed in diabetes patients who regularly take insulin injections. Several patients have been observed to develop tissue necrosis and severe infection around insulin injection sites due to insulin amyloid.^7–10^ The presence of insulin aggregates in injection sites as well as in the blood stream therefore poses a challenge for insulin to function effectively as a therapeutic^9,11^. Insulin is a peptide hormone, mainly involved in glucose homeostasis by cells, as well as cellular growth and metabolism. It is a 51 amino-acid protein that contains peptide chains A (30 amino acids) and B (21 amino acids) connected with three disulfide linkages^12,13^. Most insulin production in the human body occurs in β-cells of the pancreas and is controlled by glucose levels in the blood^14–16^. Once generated, insulin is stored within secretory granules as zinc-bound hexamers at concentrations of roughly 40 mM^17,18^. However, to become functional, insulin hexamers dissociate to monomers upon release from β-cells^18^. Depending on the local environment, such as the presence of different pH or temperature, insulin monomers can undergo partial unfolding and aggregate into amyloid species characterized by peptide strands arranged in repetitive cross β-sheet structures stabilized by hydrogen-bonds^19–23^. These amyloid aggregates are known to be inactive for cellular uptake and metabolism and have been reported to result in cellular toxicity^24–26^.

Previously, it has been reported that the conformation and structural arrangements of monomeric units within insulin fibrils contain 2, 4, and 6 protofilaments^27^. Additionally, Reif and coworkers have reported that in insulin fibrils generated under low pH, insulin monomers adopt a U-shaped conformation and fold into four β-strands in two layers within each fiber unit^28^. However, it is not clearly known whether insulin aggregates can possess more than one morphology and what key factors could give rise to structural heterogeneity within such polymorphic aggregates. Amyloid aggregates of various disease-related amyloid proteins are widely known to exhibit structural- and molecular-level polymorphism. To date, several polymorphic structures from amyloid-beta (Aβ), tau, alpha-synuclein (αS), and Amylin (IAPP) have been reported which are involved in amyloid diseases ^3,29–35^. Furthermore, several amyloidogenic proteins with well-defined folded structures have also been reported to form polymorphic fibrillar structures. Importantly, correlations between such polymorphic fibril or oligomeric structures have been established with their stability and pathological outcomes. For example, polymorphic oligomeric structures of Aβ generated in the presence of different membrane lipids are found to be differentially toxic to mammalian cells^5,36–38^. Similarly, other studies have shown that polymorphic fibrillar aggregates of αSyn exhibit prion-like properties ^39,40^. However, it is not clearly known whether insulin aggregates can possess more than one morphology and what factors could give rise to structural heterogeneity within such polymorphic aggregates.

Polymorphism within fibril structures can arise from several different phenomena: (1) environmental factors such as pH, temperature, agitation, ionic strength etc.,^41–48^ or (2) interactions of the peptides with cofactors such as other proteins, lipids, metal ions, sugars etc^49–54^. Such factors can modulate the pathways of amyloid aggregation, generating kinetically trapped intermediates or ultimate aggregates that vary in toxicity^55–57^. Previous studies suggest that lipids play a major role in inducing conformational changes and misfolding of amyloidogenic proteins like αSyn and Aβ to generate fibrillar polymorphs of distinct structures and stabilities^58–62^. Since insulin is a peptide hormone that regulates glucose uptake by the cells in different parts of the body, it is susceptible to exposure towards variable pH and ionic strength conditions to form amyloid structures such as acidic conditions in the secretory granules and neutral conditions in cytoplasm and extracellular matrix^63^. Additionally, insulin is also prone to interact with free lipids which are abundant in plasma and blood as cleavage products of triglycerides^64^. Therefore, it is important to investigate the amyloidogenic interaction of insulin with such lipids below their critical micelle concentration.

In this study we report our investigation on the effects of free lipids containing a negatively charged head group on insulin aggregation. Based on thioflavin-T (ThT) fluorescence assay, we found that the aggregation kinetics of insulin was altered by the presence of an anionic phospholipid, 1,2-Dimyristoyl-sn-glycero-3-phosphoglycerol, (DMPG) below its critical micelle concentration (CMC). Our results show that DMPG triggers the generation of low-molecular weight cytotoxic oligomers alongside longer fibrils in contrast to nontoxic insulin fibrils formed under low pH. CD, FTIR and solid-state NMR experiments show that insulin fibrils formed, along with β-sheet oligomers, in presence and absence of anionic phospholipids are β-sheet structured with a rigid core. Solid-state NMR spectra reveal a similar conformation for the fibrils grown in presence and absence of anionic and zwitterionic lipids. Importantly, the DMPG induced insulin oligomers exhibit high cellular toxicity.

## Experimental section

### Materials

Recombinant human insulin was obtained from Roche, USA (Indianapolis, IN). Dimyristoylphosphatidylglycerol (DMPG) and dimyristoylphosphatidylcholine (DMPC) lipids were purchased from Avanti Polar Lipids (Alabaster, Alabama). Sodium chloride (NaCl), sodium phosphate, and Trypsin-EDTA were purchased from Fisher Scientific (Hampton, NH). Dulbecco’s Modified Eagle Medium (DMEM) was procured from Gibco (Grand Island, NY). Copper grids used for transmission electron microscopy were supplied by Millipore Sigma (Burlington, MA), and Uranyless Stain was purchased from Electron Microscopy Sciences. Thioflavin T (ThT) was obtained from Millipore Sigma (Burlington, MA).

### ThT experiments

ThT fluorescence kinetics assays to study the self-assembly and aggregation of insulin in the presence and absence of different concentrations of DMPG were carried out on a Biotek Synergy H1 instrument with excitation at 452 nm and emission at 485 nm. Briefly, 80 µM insulin was incubated with 2, 4, 6, or 8 µM DMPG lipids or without any lipids in 10 mM sodium phosphate buffer of pH 3, with 50 µM ThT and 150 mM NaCl at 37 °C. Fluorescence was read in 96 well plates for 24-48 h under orbital shaking at 700 rpm, and with an interval of 15 minutes. Data shown has been repeated at least three times independently to ensure reproducibility.

### TEM experiments

Samples were freshly prepared by applying 10 µL of reaction samples to 300 mesh-size Formvar/Carbon Supported copper grids (Millipore Sigma-catalogue#TEM-FCF300CU) and dried for 10 minutes. Subsequently, the grids were stained with 10 µL of Uranyless stain (EMS-catalogue#22409), any excess stain was soaked on a blotting paper and the grid was dried at room temperature for 10 more minutes. Samples were imaged using a HT7800, Hitachi TEM microscope at an acceleration voltage of 100 kV. Images were collected from at least three grid regions for independent samples at magnifications between 15000x-25000x.

### CD experiments

Insulin reaction samples and resulting fibril samples were diluted accordingly to obtain a final concentration of 20 µM, and the supernatant samples were kept undiluted. CD spectra were obtained between 200-260 nm in a Chirascan instrument at room temperature. An average of three scans for each sample was plotted using OriginPro software.

### FTIR experiments

FTIR spectra were acquired using a Thermo Nicolet with diamond-ATR accessory. 1 mg of lyophilized insulin samples (insulin fibrils, oligomers and monomers) were scanned from 1500 to 1800 cm^−1^ at a resolution of 8 cm^−1^. A total accumulation of 512 spectral scans was obtained per sample, and the data were processed using a baseline correction and plotted using OriginLab8.

### Solid-state NMR experiments

Solid-state NMR experiments were performed using Bruker Avance spectrometers operating at 600 MHz (14.1 T) and 850 MHz (20.0 T) at the National High Magnetic Field Laboratory (NHMFL), Florida, and at 600 MHz (14.1 T) at Bruker facilities. The insulin fibrils (∼1 mg) were packed into a 3.2 mm rotor for experiments using a HCN triple-resonance MAS probe (NHMFL, home-built), a HX double-resonance MAS probe (NHMFL, home-built), and a HCN triple-resonance CryoProbe (Bruker). ^13^C CP-MAS and refocused-INEPT spectra were recorded under MAS spinning speeds of 12-12.5 kHz at various temperatures. Typical radiofrequency field strengths used were 68-83 kHz for ^1^H decoupling, 62.5 kHz for ^1^H hard pulses, and 50-62.5 kHz for ^13^C pulses. Experimental parameters such as cross-polarization contact time (τHC), sample temperature (T), recycle delay (d1), number of scans (NS), and *J*-evolution times are provided in the figure legend. ^13^C chemical shifts were externally referenced to adamantane’s CH2 peak at 38.48 ppm with 0 ppm for ^13^C peak of tetramethylsilane (TMS).

### Solution-state NMR experiments

Solution NMR experiments were performed on a 700 MHz Bruker NMR spectrometer equipped with a 5 mm-TCI cryoprobe. NMR samples were prepared and kept on ice before being loaded to NMR probe for data acquisition. ^1^H NMR data was acquired with 1024 scans, 1.25 s recycle delay, and water suppression by the WATERGATE sequence. Spectra were collected at 35 °C, and the sample was kept in the magnet at the same temperature between data acquisitions at different timepoints.

### Cellular cytotoxicity

NIH3T3 fibroblasts were maintained in Dulbecco’s Modified Eagle Medium (DMEM) supplemented with 10% fetal bovine serum (FBS) and 1% penicillin-streptomycin. The cells were maintained at 37 °C in a humidified incubator with 5% CO2. For subculturing, when the cells reached 70-80% confluency, the culture medium was removed, and the cells were washed with phosphate-buffered saline (PBS). 0.25% trypsin-EDTA solution was added to detach the cells, which were centrifuged to obtain a pellet. This pellet was resuspended in fresh culture medium and transferred to new culture vessels to ensure optimal growth conditions for experimental purposes. NIH3T3 cells (1×10□J) were seeded in 96-well plates and treated with different aggregate species of insulin for 36 hours. After treatment, cellular viability was assessed using the CCK-8 assay and absorbance was measured at 450 nm to evaluate cell viability.

## Results and discussion

### Anionic phospholipids below their critical micelle concentration augment insulin aggregation

To investigate the effect of negatively charged phospholipids below their critical micelle concentration on amyloid formation by insulin monomers, we performed ThT fluorescence assays of 80 µM insulin incubated with 2-8 µM DMPG phospholipids, which is below the reported CMC of DMPG (i.e. 11µM)^65^ in phosphate buffer at pH 3.0 and 37 °C with an orbital shaking at 700 rpm (Figure 1a). We found that the presence of DMPG accelerated insulin aggregation drastically, shortening the lag time to 1-2 hours after the start of the reactions whereas the insulin aggregation exhibited a lag time up to 10-12 hours in the absence of lipids. While these results support previous findings on the acceleration of insulin aggregation in presence of anionic lipids^66^, we found that, within the lipid concentration range used here, the ThT fluorescence kinetics were independent of the concentration of lipids in the sample. Additionally, we screened the effect of zwitterionic DMPC (1,2-Dimyristoyl-sn-glycero-3-phosphocholine) lipids across various concentrations near its CMC (6 nM)^65^. We found that DMPC has negligible effect on insulin at and below the CMC, however, it delays insulin aggregation up to 4h above CMC at 8nM (Supplemental Figure S3). We further performed a secondary structure analysis of the bulk solution at 16h after the start of the aggregation reaction with CD spectroscopy. We observed that by 16h, the alpha helical structure of insulin at t=0 h, as indicated by the double-minima at 222 and 210 nm, had transitioned to β-sheet structure, which was indicated with a minimum at 218 nm^67^.

**Figure 1.**
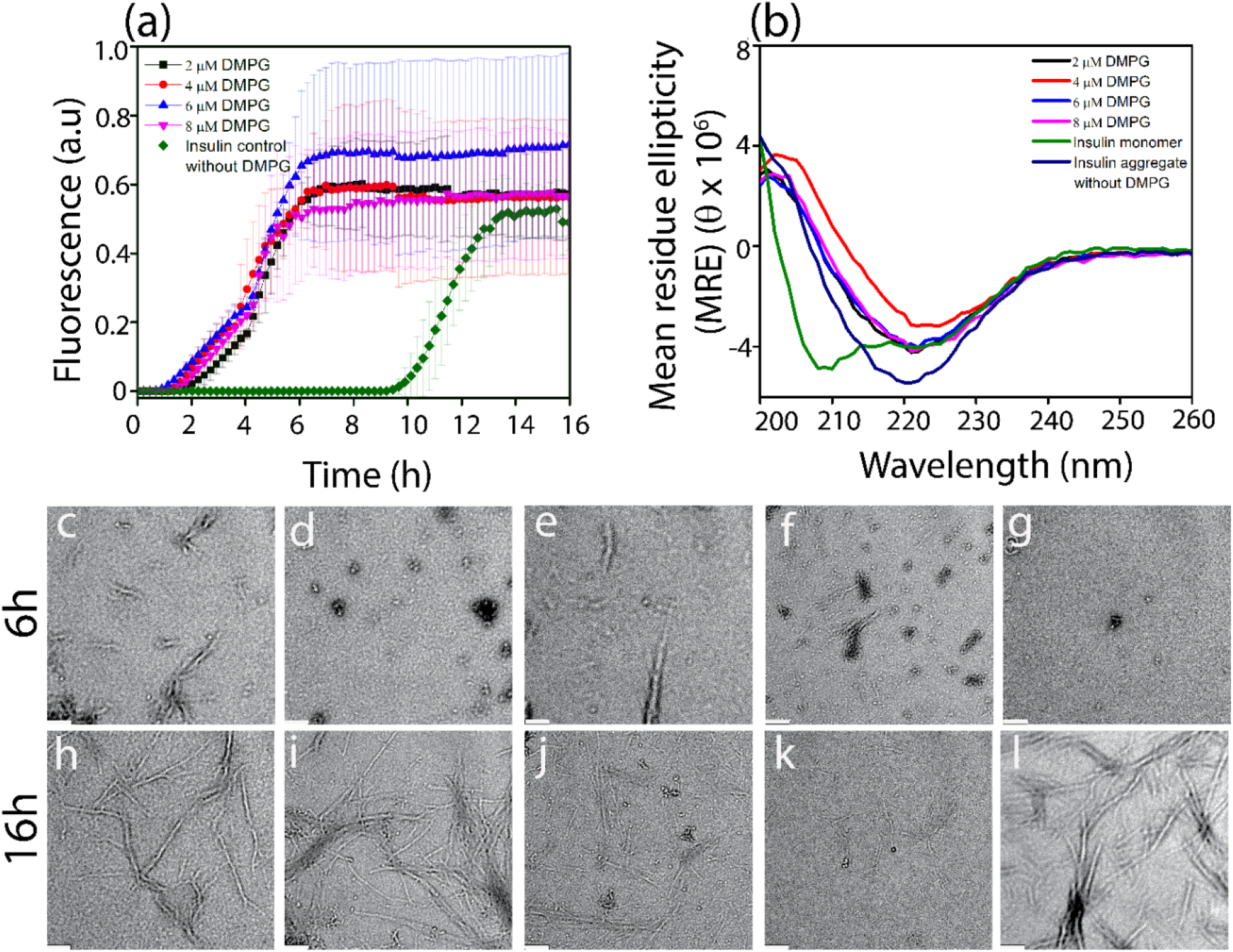
ThT fluorescence kinetics (a) and circular dichroism (CD) spectra (b) of 80 µM insulin with 0 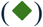, 2 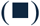, 4 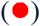, 6 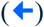, and 8 µM 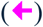 DMPG. Insulin and DMPG were incubated in 10 mM sodium phosphate buffer (pH 3, 150 mM NaCl, 50 µM ThT) at 37 °C and with agitation at 700 rpm while insulin monomers were freshly prepared in 10 mM sodium phosphate buffer at pH 3. (c-l) TEM images of insulin aggregates generated in the ThT fluorescence assay with 2(c, h), 4(d, i), 6(e, j), 8 (f, k) µM DMPG and without DMPG (g, l); these aggregates were collected after 6h (c-g) or 16h (h-l). The indicated scale bar is 200 nm.

Previous work has demonstrated that amyloid aggregation and formation of fibrils are preceded by the generation of intermediate oligomeric structures^37,38,40,56^. The structure, conformation and stability of such oligomers are dependent upon environmental factors such as temperature, pH, ionic strength of the buffer, or the presence of other interacting molecules such as proteins and lipids^68,69^. Therefore, to investigate whether the presence of DMPG lipids affects the generation of intermediate species along the aggregation pathway of insulin, we performed transmission electron microscopy (TEM) imaging on insulin aggregates generated with various concentrations of DMPG (Figure 1 c-l). Samples were collected after incubation at 37 °C and under continuous shaking at 700 rpm for 6 and 16 hours. We observed that with 2 µM DMPG, insulin formed short fibrils within 6 h of incubation (Figure 1c) that became larger and denser by 16h (Figure 1h). However, when the concentration of DMPG was increased to 4, 6 and 8 µM, a mixture of small spherical oligomeric intermediates and short fibrils of insulin emerged within 6 h of incubation (Figure 1d-f). While the intermediates largely transitioned to larger fibrils by 16 h in the case of 4 µM DMPG (Figure 1i), the oligomeric intermediates persisted alongside larger fibrils after 16h for insulin samples with 6 µM and 8 µM DMPG (Figure 1j, k). Insulin aggregated without lipids or with DMPC lipids resulted in the formation of a small population of spherical intermediates at 6 h (Supplemental Figure S3(b-f) & Figure 1g) that completely turned into larger fibrils within 16 h (Supplemental Figure S3(g-k) & Figure 1l). Therefore, our findings suggest that insulin aggregates via an alternative pathway by forming distinct oligomeric species in the presence of certain concentrations of anionic phospholipids. We also attempted to investigate the formation of different intermediate species upon insulin aggregation with solution NMR as done previously in other studies related to amyloid aggregation^70,71^. We observed similar peaks in ^1^H spectra at 0 h and at 24 h for insulin samples incubated at 35 °C with or without DMPG but without shaking the sample in the NMR tube (Supplemental Figure S2). The absence of peak broadening or intensity loss is indicative of negligible or delayed aggregation^72^. These observations emphasize the significance of agitation for insulin aggregation.

### Small soluble cytotoxic oligomers are generated alongside larger fibrils

TEM images of insulin aggregates generated in presence of 6 µM and 8 µM DMPG indicate that a mixture of small oligomers and fibrils was present in the reaction sample and stable for at least 16 h (Figures 1j, k). When temperature was increased to speed up the aggregation process, we observed that the oligomers were still generated alongside fibrils. For example, incubation of 6 µM DMPG and insulin as previously except at a higher temperature (70 °C) resulted in a mixture of oligomers and fibrils in the presence of DMPG (Figure 2g) but not in its absence (Figure 2j). Next, we isolated the insoluble fibril fraction from the soluble oligomeric fraction using high-speed centrifugation. The samples containing the insoluble fibril pellet formed with or without DMPG displayed a large hydrodynamic diameter greater than 1000 nm (Figures 2a, b), as observed by DLS. Another size distribution peak was also observed near the smaller diameter region at 1-10 nm, which may be due to dissociated monomeric or oligomeric species. The presence of large fibrils in these samples was further confirmed by TEM (Figures 2h, k). DLS profiles of the soluble fractions revealed the presence of oligomeric structures of about 10-15 nm for insulin incubated with DMPG (Figure 2d), whereas species with diameter of 2-4 nm corresponding to insulin monomers (Supplemental Figure S1) were observed in the soluble fraction of insulin incubated without DMPG (Figure 2e). Accordingly, small spherical structures were visible under TEM for the soluble fraction of insulin incubated with DMPG (Figure 2i). This further confirms the presence of oligomers. In contrast, negligible aggregate species were visible in the soluble fraction of insulin aggregates formed without lipids. The secondary structure of both pellet and supernatant for each sample was further analyzed using CD spectrometry. It was observed that all fibril samples showed the presence of twisted β-sheet structure evident from a minimum near 230 nm^73^ as opposed to insulin monomers which mostly had α-helical structure evident from the double minima near 210 and 222 nm (Figures 2c and f). Furthermore, the presence of the minimum at 218 for the soluble fraction of insulin without DMPG indicated the presence of a small population of aggregates with β-sheet structure even after centrifugal separation of the fibrillar pellet. The presence of β-sheet in both insulin fibrils with and without DMPG was confirmed by the peaks near 1630 cm^-1^ in FTIR spectra (Figure 2n). However, the oligomeric fraction of DMPG and insulin displayed a peak at 1670 cm^-1^ which is attributed to β-turns^74,75^. In contrast, FTIR spectrum of insulin monomers displayed a peak at 1647 cm^-1^, indicating a mixture of α-helical and disordered conformations.

**Figure 2.**
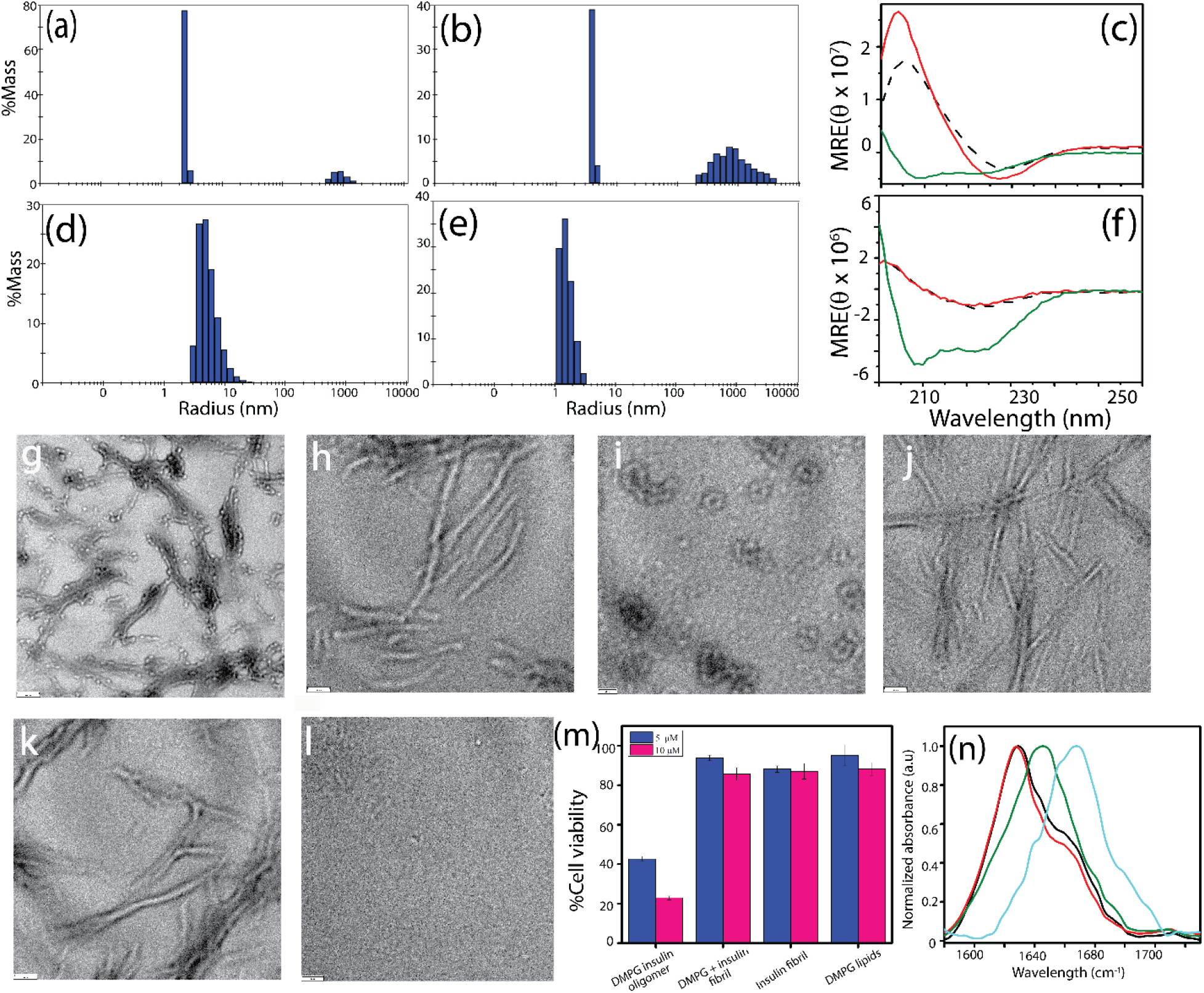
DLS profiles (a,b,d,e) and CD spectra (c,f) of pellet (a,b,c) and supernatant (d,e,f) fractions of 80 µM insulin monomer reaction with (a, d in DLS; black trace in CD) and without (b, e in DLS; red trace in CD) 6 µM DMPG centrifuged at 18000x g for 30 minutes after incubation at 70 °C under continuous agitation at 700 rpm for 16 h. CD spectra of insulin monomers at 0 h (green traces in c and f). TEM images of total (g, j), pellet (h, k), and supernatant (i, l) fractions of 80 µM insulin monomer reaction with (g, h, i) and without (j, k, l) 6 µM DMPG. The scale bar is 100 nm. (m) CCK-8 cytotoxicity assay on NIH3T3 cells for 5 and 10 µM insulin oligomers catalyzed by DMPG, and insulin fibrils generated with and without DMPG, and DMPG. (n) FTIR spectra of insulin fibrils generated with (□) and without 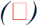 DMPG lipids, insulin oligomers generated with DMPG lipids 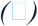, and insulin monomers 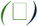.

To elucidate the structural features of insulin fibrils, magic angle spinning (MAS) solid-state NMR experiments were conducted on fibrils. The natural-abundance ^13^C CP-MAS NMR spectra of insulin fibrils, prepared in the absence and presence of DMPG and DMPC lipids, were found to be similar (Figures 3a-c). These results indicated that the backbone conformation of insulin fibrils were not altered by the presence of lipids (DMPG or DMPC), as evidenced by the similar ^13^Cα and ^13^CO chemical shifts observed for all three samples. These findings are in good agreement with the observations from FTIR spectroscopy. To investigate the side-chain environments of the insulin fibrils, refocused-INEPT experiments were utilized. The observed peaks in refocused-INEPT spectra were predominantly attributed to fibril side chains since the fibrils generated under free lipids (6 *μ*M DMPG or 6 nM DMPC) were expected to contain negligible lipids due to discarding the lipid containing soluble fraction via centrifugation prior to NMR measurements. A comparison of refocused-INEPT spectra revealed notable differences between the fibrils formed in the presence of DMPG and DMPC (Figure 3d-f). Specifically, the DMPC-catalyzed fibrils exhibited distinct aliphatic and aromatic signals (Figure 3d,e), suggesting subtle structural variations in the fibrils influenced by the lipid environment. These findings suggest that the fibrils formed in the presence of DMPG and DMPC likely to exhibit distinct structural arrangements. These observations indicate that the lipid environment influences the fibril structure and potentially alters the conformational flexibility or intermolecular interactions. This variation underscores the impact of lipid-based interactions on the kinetics of fibril formation and structure of insulin aggregates. In order to probe the interaction between monomeric insulin and lipids, ^1^H NMR spectra of monomers of insulin in the presence of DMPG were acquired for 24 hours (Supplemental Figure S2). However, no measurable changes in the spectra were observed indicating no aggregation of insulin monomers. This further confirms the need for mechanical agitation to induce the aggregation of insulin monomers. On the other hand, solution NMR spectra (Supplemental Figure S6) of insulin samples at different time points of aggregation under agitation with or without lipids were found to be significantly different from that of insulin monomers (Supplemental Figure S2). These results suggest that insulin quickly aggregates (at least within 6 hours) that are large in size for detection by solution NMR and therefore do not provide narrow ^1^H spectral lines in solution NMR spectra. The observed peaks in ^1^H NMR spectra of agitated insulin samples (with or without lipids) likely to be from monomers present in the sample. Additional solid-state and solution NMR experiments using isotopically (^13^C) enriched insulin samples could b useful to obtain piercing insights into lipid-insulin interactions.

**Figure 3.**
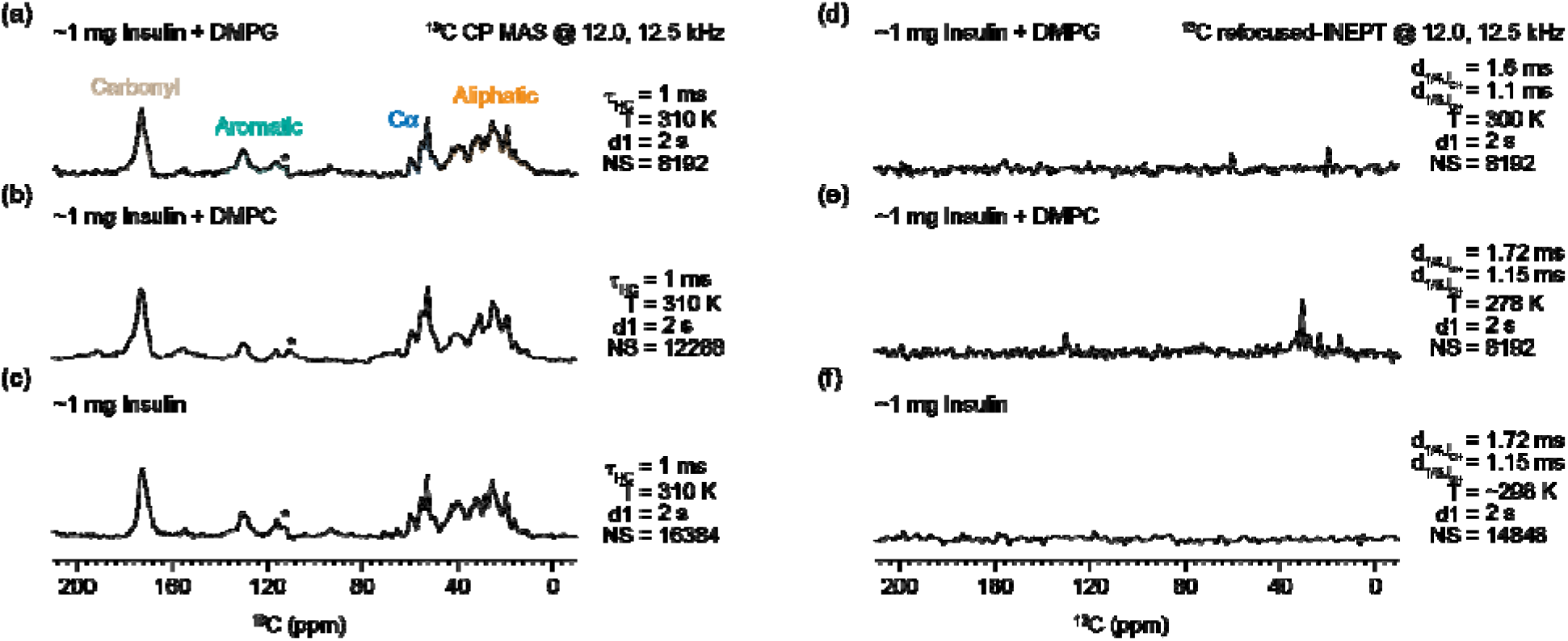
Natural-abundance ^13^C MAS NMR spectra of insulin aggregates prepared in presence of DMPG and DMPC (a,b,d,e) and without lipids (c,f). The CP-MAS spectra were obtained at 12 kHz (a,c,d) and at 12.5 kHz (b,e,f) at the indicated contact-time (*τ*HC) for cross-polarization, temperature (T), recycle delay (d1) and number of scans (NS). ^13^C MAS NMR spectra obtained using refocused-INEPT showed no peaks from insulin fibers generated without lipids (f) whereas a few peaks were observed for the insulin-lipids samples (d,e) which are likely to be from aromatic and aliphatic side chains of insulin as the samples had very little amount of lipids. All spectra were obtained on a 600 MHz Bruker NMR spectrometer except the refocused-INEPT spectra (j and k) that were acquired using an 850 MHz Bruker NMR spectrometer. Additional CPMAS NMR spectra are given in Supplemental Figure S7.

Previous research by several groups has established that amyloid aggregates of the same protein generated under different conditions display variable toxicity towards mammalian cells. Therefore, we analyzed the cytotoxic effects of different aggregate species of insulin towards NIH3T3 fibroblast cells. Fibril pellets generated with and without DMPG lipids and oligomers generated with DMPG lipids were incubated with NIH3T3 cells for 36h. Then, a CCK-8 assay was carried out to assess the dehydrogenase inactivation in cells as a measure of cellular viability. Negligible toxicity was observed in the case of fibril pellets of insulin samples with and without DMPG. Interestingly, the soluble fraction from insulin-DMPG samples containing the insulin oligomers showed the highest toxicity where cellular viability decreased to 40% and 20% with 5 and 10 µM insulin concentrations, respectively.

## Discussion

Our study underscores the impact of the anionic phospholipid DMPG on insulin aggregation, showing distinct aggregation pathways that differ from those observed under low pH conditions without lipids. The presence of negatively charged phospholipids has been shown to accelerate the aggregation of insulin in our study as well as in previous studies^76^. These findings suggest that the negative charge of lipids plays a crucial role in lipid-protein interactions and in the self-assembly process leading to protein aggregation. The interaction between the negatively charged lipids and insulin can be explained by electrostatic interactions. Since insulin has a pI of 5.3, the net charge on the insulin in acidic pH is positive. Therefore, electrostatic interaction between the positively charged insulin and negatively charged DMPG is likely to facilitate insulin-insulin interactions, promoting the self-assembly process to form the oligomers and fibrils (Supplemental Figure S5). On the other hand, minimal interaction was observed when insulin was incubated with DMPC, which contains a zwitterionic headgroup, below its CMC. Insulin aggregation was somewhat attenuated in the presence of DMPC above its CMC. Interestingly, it is also noted that a change in the rate of aggregation corresponded with changes in the secondary structure of insulin. While incubation of insulin with DMPG led to the formation of small spherical oligomers and larger fibrils, incubation with DMPC resulted in a mixture of shorter and larger fibrillar structures (Supplemental figure S4).

Based on the experimental results reported in this study, we propose that under certain concentrations of DMPG, insulin aggregates through a different aggregation pathway due to the presence of oligomeric aggregates. It is possible that the decrease in the lag-time also correlates to this change. Several other reports have shown that lipids can act as nucleation sites for amyloid aggregation, and our results are consistent with reports that emphasize the importance of lipid charge and composition in dictating the aggregation routes^76,77^. Specifically, studies on amyloid β (Aβ) and α-synuclein have shown that anionic lipids like DMPG promote protein-lipid interactions that lead to oligomeric intermediates, like what we have observed with insulin^78,79^. These intermediates are considered to be crucial participants in amyloid toxicity, as observed in this study, and their formation may depend on lipid-protein electrostatic interactions. Furthermore, the effect of lipids on aggregation kinetics observed in our ThT experiments supports previous research demonstrating that the presence of lipids can accelerate or alter the trajectory of protein aggregation. This shift in the aggregation pathway away from pure fibril formation to a combination of oligomers and fibrils has been noted in studies on transthyretin and prion proteins, where lipid environments lead to differences in aggregation patterns^80–82^. These findings add to the growing body of research showing that lipids can act as modulators of amyloid fibrillation in various protein systems, including insulin, and contribute new insights into the structural and cytotoxic characteristics of lipid-associated insulin oligomers. Our TEM images, combined with DLS data, clearly showed the presence of both fibrils and oligomeric intermediates in insulin-DMPG mixtures, whereas insulin alone predominantly formed fibrils without discernible oligomers at later stages of aggregation. This observation is corroborated by literature reports, where lipid-induced stabilization of small oligomeric species has been identified as a common phenomenon in amyloid systems. For instance, recent work on IAPP (islet amyloid polypeptide) demonstrated that lipids can stabilize early-stage oligomers that later develop into fibrils^83^. The small hydrodynamic radii (around 15 nm) of the oligomers observed in our study are similar to sizes reported for toxic oligomers in other amyloid proteins, such as Aβ and tau^84,85^. This similarity suggests that the formation of lipid-bound oligomers may be a general mechanism by which lipids influence the amyloid aggregation of different proteins.

Furthermore, a shift in secondary structures is also observed for insulin while transitioning from monomeric to aggregated structures. The shift from α-helical structures in native insulin to β-sheet-rich structures during aggregation was confirmed by CD spectroscopy, consistent with other amyloid-forming proteins, where β-sheet formation is critical for fibrillation. Our results echo studies showing that amyloidogenic proteins tend to adopt β-sheet-rich conformations when transitioning to aggregated states in lipid environments^86,87^. The FTIR data, showing β-turns in the oligomers (carbonyl stretch at 1660 cm□^1^), suggest that these intermediates are structurally distinct from both native and mature fibrils of insulin. Previous studies using FTIR on Aβ oligomers also noted similar spectral features, indicating β-turn structures that contribute to their unique properties and toxicity^58^. These findings reinforce the idea that lipid-induced oligomers may have distinct folding patterns, which could contribute to their stability and toxicity. Insulin oligomers were also shown to be the most cytotoxic aggregate species in our study. This toxicity is believed to be correlated with oligomer structure of amyloids in which hydrophobic surfaces are exposed in β-sheets^56,88–90^. This hydrophobic quality can allow oligomers to infiltrate cell membranes and cause cell death. Several studies focused on Alzheimer’s disease have shown that small soluble oligomers of Aβ are far more cytotoxic than fibrils due to their ability to disrupt cellular membranes and interfere with cellular processes^36,37,91^. Similarly, α-synuclein oligomers have been shown to be toxic to neuronal cells through mechanisms that likely involve membrane disruption^92–94^. Our finding that insulin oligomers in the presence of DMPG lipids are more toxic to fibroblasts further supports the hypothesis that lipid-induced oligomers may share similar toxicity mechanisms across different amyloid proteins. The increased toxicity of lipid-associated insulin oligomers in NIH3T3 fibroblast cells observed in our study is consistent with a wealth of literature suggesting that oligomeric intermediates, rather than fibrils, are the primary toxic species in amyloid-related diseases. However, the specific lipid-driven mechanism underlying the cytotoxicity of these intermediates remains unclear, and it is uncertain which part of the insulin molecule (A or B chain) interacts with DMPG to promote toxicity. Similar questions have arisen in other studies, such as those on Aβ, where the exact nature of lipid-protein interactions contributing to toxicity is still under investigation. Future structural studies, possibly using NMR and cryoEM, could provide more detailed insights into these specific interactions.

Our study adds to the growing understanding of how lipid environments influence protein aggregation and toxicity. The interaction of insulin with DMPG leading to the formation of toxic oligomers has broad implications for amyloid-related diseases, especially those where insulin aggregation is implicated. Moreover, this research could have therapeutic implications in designing inhibitors that target specific lipid-protein interactions to prevent toxic oligomer formation. However, it remains to be investigated whether these oligomers further seed insulin monomers to form structurally distinct fibrils of insulin. Furthermore, since insulin and its bioavailability in its native form is involved in diabetes, it is crucial to understand whether insulin oligomers affect cross-seeding of IAPP. Moreover, insulin’s ubiquitous presence in various organs also makes it imperative to understand its crosstalk with other amyloidogenic proteins such as Aβ and αSyn. Future studies should focus on identifying the specific molecular interactions between insulin and DMPG to uncover which regions of the insulin molecule are responsible for lipid-induced aggregation. Additionally, understanding how these lipid-associated oligomers exert their toxic effects on cells could pave the way for novel therapeutic strategies aimed at mitigating amyloid toxicity in both diabetes and other amyloid-associated conditions. In conclusion, our findings demonstrate that anionic lipids such as DMPG significantly modulate insulin aggregation, leading to the formation of small, toxic oligomers that differ structurally from both monomers and fibrils. These results are consistent with existing studies on other amyloid proteins and suggest that lipid-induced oligomers may represent a key target for understanding and treating amyloid diseases.

## Supporting Information

Hydrodynamic radius determination for Insulin monomers at pH 3 using DLS data, ^1^H NMR spectra of insulin monomers in the presence of DMPG, ThT fluorescence kinetics and TEM of insulin with DMPC, TEM images (of total, pellet, and supernatant fractions insulin monomer reaction with DMPC), structure of insulin and DMPG and DMPC lipids, ^1^H NMR spectra of insulin aggregates generated with and without DMPG at 6h or 24h, natural-abundance ^13^C MAS NMR spectra of insulin aggregates prepared in the presence of DMPG or DMPC and without lipids

## Author Contribution

JS and AR conceived and planned the project. JS, AW, MCDW, SM, RF, and NK carried out the experiments. JS, MCDW, SM and AR interpreted the results. JS prepared the first draft of the manuscript, and SM and AR edited and revised the manuscript. JA and AR supervised the research. All authors approved the final version of the manuscript. AR obtained funding and directed the project.

## Acknowledgements

We thank Dr. Jochem Struppe, Dr. Barbara Perrone and Alia Hassan from Bruker for acquiring one of the CPMAS NMR spectra for this study.

