## Supporting Information for "Anionic lipid catalyzes the generation of cytotoxic insulin oligomers"

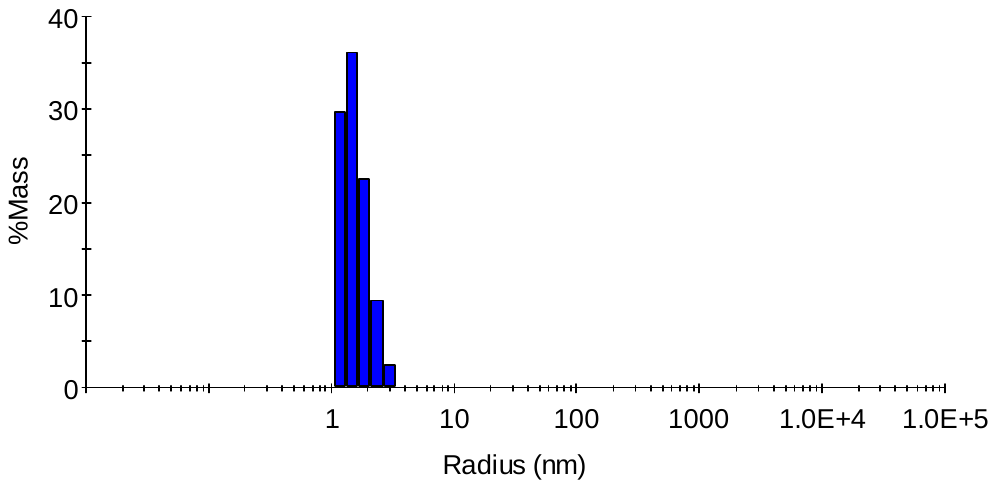
**Supplementary figure1**

*Figure S1: Dynamic light scattering (DLS) profile of freshly prepared insulin monomers (10 µM) in 10 mM phosphate buffer pH 3.*

**Supplementary figure 2**

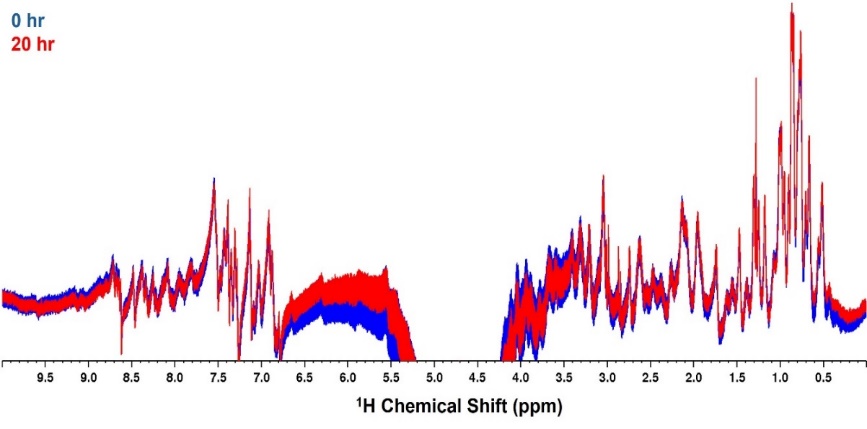

*Figure S2: ^1^H NMR spectra of insulin monomers in the presence of DMPG at 37 °C in 10 mM sodium phosphate pH 3 without any shaking from 0 to 24 h acquired on a 700 MHz NMR spectrometer using a cryoprobe.*

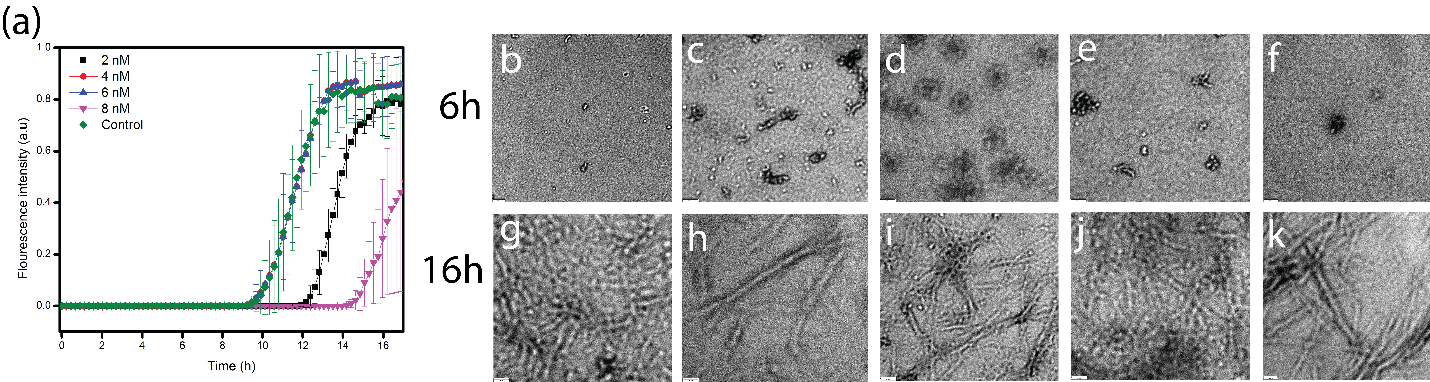
**Supplementary figure 3**

*Figure S3: ThT fluorescence kinetics (a) of 80 µM insulin with 0 (⯁), 2 (◼), 4 (⚫), 6(□), and 8 µM (□) DMPC. Insulin\and DMPG were incubated in 10 mM sodium phosphate buffer (pH 3, 150 mM NaCl, 50 µM ThT) at 37 °C and with agitation at 700 rpm while insulin monomers were freshly prepared in 10 mM sodium phosphate buffer at pH 3. (b-k) TEM images of insulin aggregates generated in the ThT fluorescence assay with 2(b, g), 4(c, h), 6(d, i), 8 (e, j) µM DMPG and without DMPG (f, k); these aggregates were collected after 6h (c-g) and 16h (h-l). Scale bar is 200 nm.*

**Supplementary figure 4**

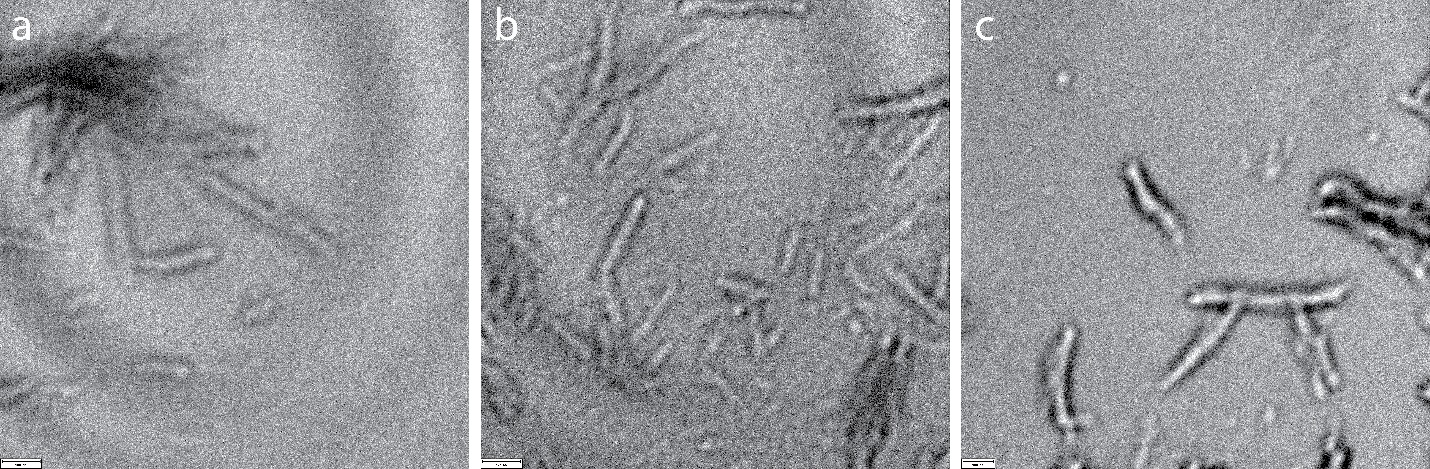

*Figure S4: TEM images of total (a), pellet (b), and supernatant (c) fractions of 80 µM insulin monomer reaction with 6 nM DMPC. The scale bar is 100 nm.*

**Supplementary figure 5**

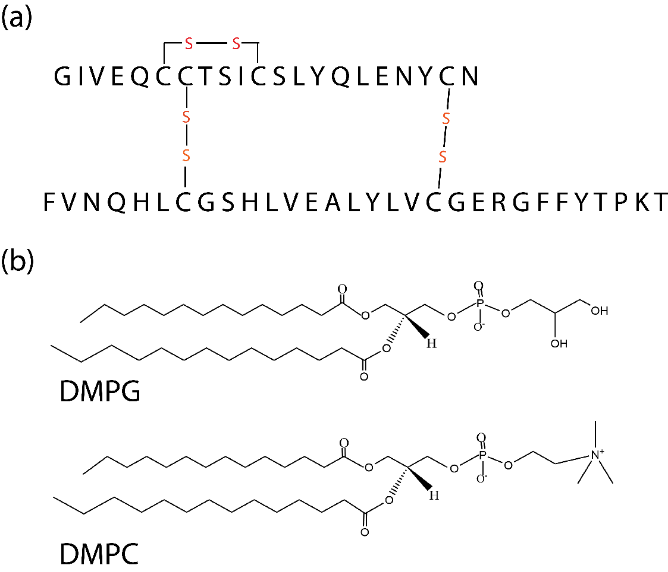

*Figure S5: Molecular structure of Insulin and DMPG and DMPC lipids used in this study.*

**Supplementary figure 6**

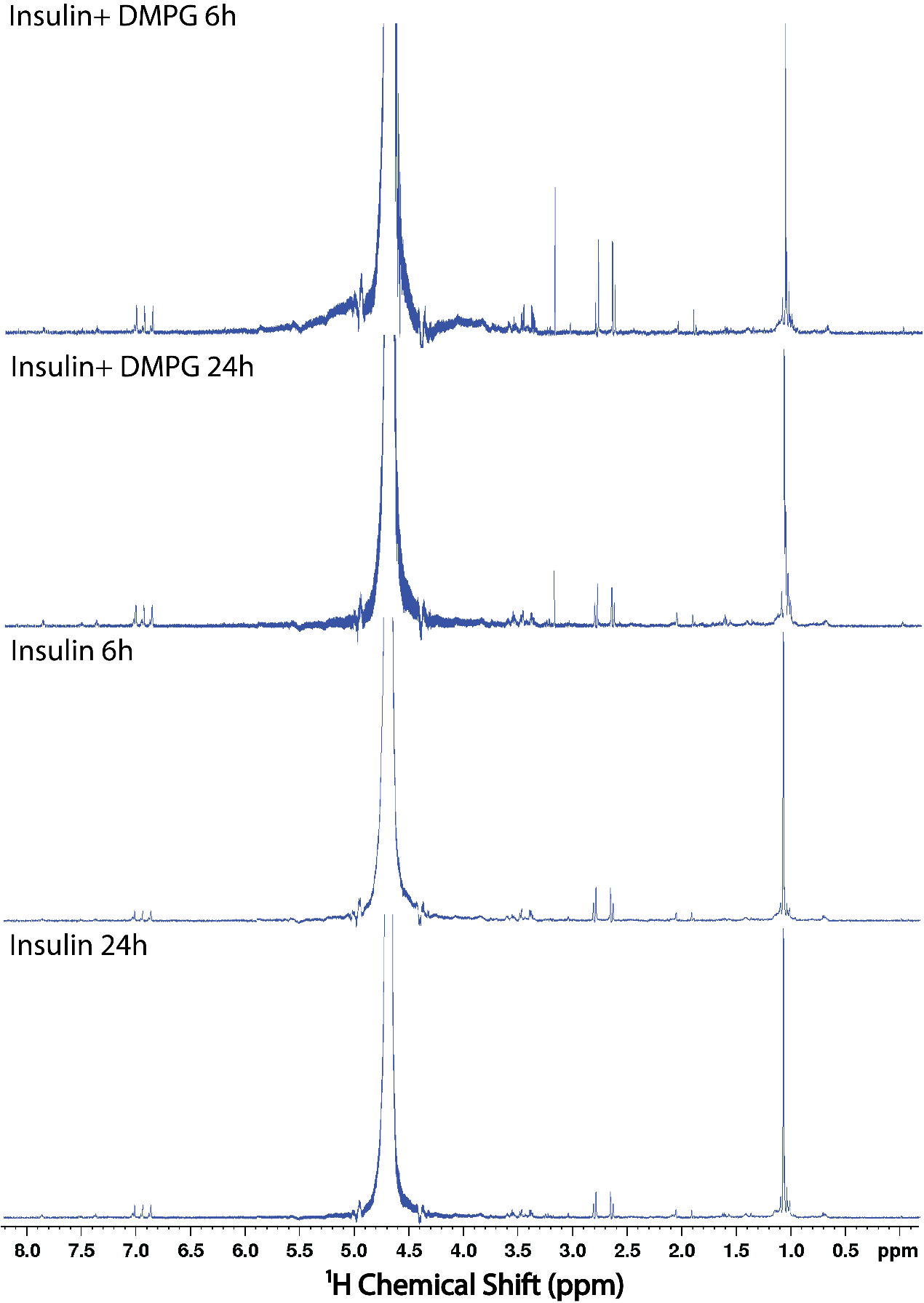

*Figure S6: ^1^H NMR spectra of insulin aggregates generated with and without DMPG at 6h or 24h upon shaking at 700 rpm at 37 °C in 10 mM sodium phosphate pH 3. Spectra were acquired on a 700 MHz NMR spectrometer using a cryoprobe.*

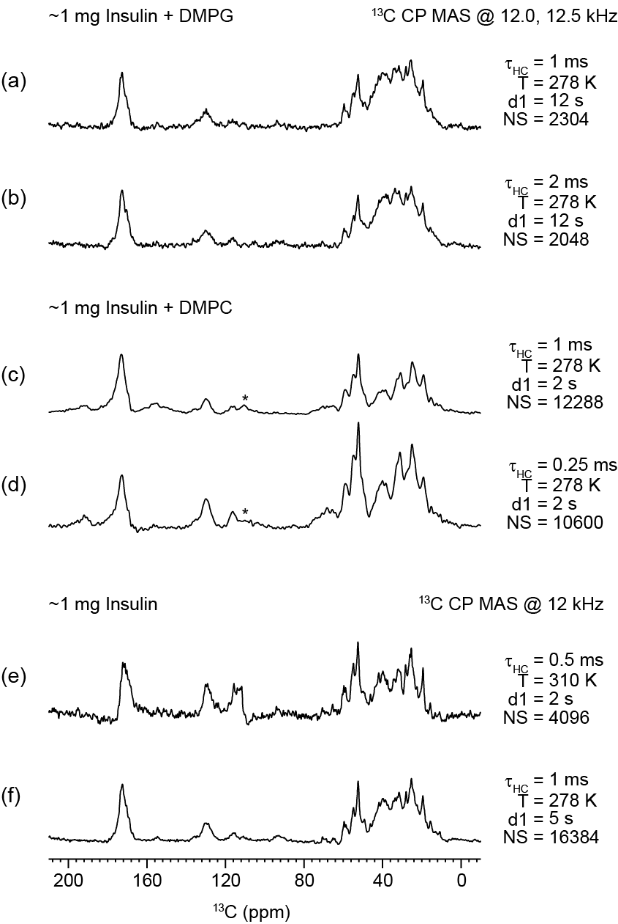

*Figure* *S7.* Natural-abundance ^13^C MAS NMR spectra of insulin aggregates prepared in presence of DMPG or DMPC (a,b,d,e) and without lipids (c,f). Other NMR experimental parameters used to acquire these spectra are as mentioned in Figure 3.
